# Behavioral and Neural Representations en route to Intuitive Action Understanding

**DOI:** 10.1101/2021.04.08.438996

**Authors:** Leyla Tarhan, Julian De Freitas, Talia Konkle

## Abstract

When we observe another person’s actions, we process many kinds of information – from how their body moves to the intention behind their movements. What kinds of information underlie our intuitive understanding about how similar actions are to each other? To address this question, we measured the intuitive similarities among a large set of everyday action videos using multi-arrangement experiments, then used a modeling approach to predict this intuitive similarity space along three hypothesized properties. We found that similarity in the actors’ inferred goals predicted the intuitive similarity judgments the best, followed by similarity in the actors’ movements, with little contribution from the videos’ visual appearance. In opportunistic fMRI analyses assessing brain-behavior correlations, we found suggestive evidence for an action processing hierarchy, in which these three kinds of action similarities are reflected in the structure of brain responses along a posterior-to-anterior gradient on the lateral surface of the visual cortex. Altogether, this work joins existing literature suggesting that humans are naturally tuned to process others’ intentions, and that the visuo-motor cortex computes the perceptual precursors of the higher-level representations over which intuitive action perception operates.

## INTRODUCTION

Watching other people’s actions is a major component of natural vision. These actions make up a rich and varied domain of visual input — in a typical day, we might see a child building a snowman, someone re-stocking shelves at a grocery store, or a runner jogging through a park. We not only see these actions, but also understand them — we can infer at a glance how experienced the runner is, and that her goal is to exercise. Underlying this capacity is a series of social-visual computations, from higher-level inferences about an actor’s mental state (Dodell-Feder et al., 2011; Koster-Hale et al., 2017; Samson et al., 2004), to the intermediate-level perceptual representations of individual body parts, objects, and physical properties like force and momentum (Downing et al., 2001; Rosch et al., 1976; Singh, 2015; Fischer et al., 2016; Schwettmann et al., 2019; Tarhan and Konkle, 2020b). Further, all of these computations are initially embedded in early sensory representations, which capture lower-level properties like edge orientations and motion direction across the visual field (Giese and Poggio, 2003; Hubel and Wiesel, 1962).

Yet, not all of these social-visual computations necessarily influence our intuitive perceptions of actions — the things that humans naturally notice about actions and that inform our behavior (Vallacher and Wegner, 1989). For example, humans can intuitively distinguish between a person who is jogging for fitness and a person who is running to catch a bus. But, many of the properties that the visual system processes during action perception —-such as edge orientation —-may not influence this level of perception. Thus, our question is: what properties underlie this intuitive understanding of actions and what makes them similar or different from each otherã

The actors’ intentions are one property that may be tied to our intuitions about action similarity. There is a rich developmental and social psychology literature demonstrating that actors’ intentions are key to our understanding of actions, and that humans naturally process others’ mental lives and goals when watching their actions. For example, infants expect others to reach for valuable objects in the most efficient way possible and are sensitive to whether an action’s goal was completed (Gergely and Csibra, 2003; Jara-Ettinger et al., 2016; Liu et al., 2017; Reid et al., 2007; Schachner and Carey, 2013). As adults, we also naturally describe actions in terms of their goals, suggesting that they are a particularly salient property —-we say that someone “gave money to a homeless person,” rather than that they “grasped a dollar and extended it to a homeless person” (Spunt et al., 2011). In addition, we attribute motives and agency even to simple shapes that move in a way that indicates animacy (Heider and Simmel, 1944; see also De Freitas and Alvarez, 2018; Isik et al., 2018). Finally, neuroimaging work suggests that we naturally represent other people in terms of the mental states that they habitually experience (Thornton et al., 2019a). Regions of the brain that process social information even seem to automatically predict others’ future mental states (Thornton et al., 2019b) and the next event in a narrative (Richardson and Saxe, 2020). All of this research suggests that similarities in actors’ goals and intentions may influence intuitive perceptions of action similarity, because we naturally process these elements of an action when we see it.

In addition to these inferences about the actor’s mental state, it is possible that directly perceptible properties also influence intuitive action similarity. For example, perhaps running and walking seem similar because they involve similar leg movements, or because they both tend to occur outdoors. Recent work on the visual processing of actions has identified several such intermediate-level visual features that might influence intuitive action perception. These include an action’s movement kinematics, such as the body parts involved in an action and the movement’s speed and direction (e.g., Pitcher and Ungerleider, 2020; Tarhan and Konkle, 2020b); the people, objects, and spaces that actions are directed at (Tarhan and Konkle, 2020b); and the general configuration of an actor’s body relative to another person (Abassi and Papeo, 2020; Isik et al., 2017; Papeo et al., 2017). These properties might influence intuitive action perception because they are useful for inferring an action’s meaning; for example, body position can signal whether two actors are interacting (Isik et al., 2017; Papeo et al., 2017) and other kinds of motion features may influence moral judgments like blameworthiness (De Freitas and Alvarez, 2018).

Finally, it is possible that even the basic visual appearance of an action scene also influences our intuitions about action similarity. Increasingly, research on object and scene perception finds that low- and mid-level visual features can influence higher-order perceptual processing (Greene and Hansen, 2020; Groen et al., 2018; Long et al., 2018; Oliva and Torralba, 2006). For example, curvature features can influence perceptions of real-world size (Long et al., 2018) and distributions of spatial orientations differ between indoor and outdoor scenes (Oliva and Torralba, 2006). However, these early levels of representation may only be useful in initial stages of action analyses without entering into our intuitive understanding of actions.

In the present work, we probe how these different levels of representation contribute to intuitive action understanding. To do so, we used both behavioral and neuroimaging analyses to explore the nature of intuitive action representations. To measure the intuitive similarities between a set of short action videos, we used a multi-arrangement task, in which participants arranged videos according to their intuitive similarity (Kriegeskorte and Mur, 2012). This task has successfully been used to study object and scene similarity (e.g., Jozwik et al., 2016; Groen et al., 2018). These action stimuli depicted everyday sequences of movements – such as chopping vegetables – in naturalistic, 2.5-second videos (from Tarhan and Konkle, 2020b). Importantly, these videos include the contextual information derived from the scene and the action’s effects on the surrounding people, objects, and scene, and not just the actor’s isolated movements (e.g., Haxby et al., 2020; Tucciarelli et al., 2019). These naturalistic and representatively-sampled stimuli contrast with targeted approaches that use abstract stimuli like verbs (e.g., Bedny and Caramazza, 2011) or tightly controlled videos of a small set of actions (e.g., Wurm et al., 2017). This approach makes it easier to draw general conclusions about natural action perception (Haxby et al., 2020).

Next, we operationalized hypotheses drawn from the developmental, social, and vision literatures by collecting human judgments of the action videos’ similarity along three broad dimensions: the actors’ goals, the actors’ movements, and the videos’ visual appearance. We then assessed each of these hypotheses, using predictive modeling techniques that have been used to study intuitive object and scene perception (e.g., Jozwik et al., 2016; Groen et al., 2018). Finally, we mapped regions of the visuo-motor cortex that respond according to these different representational formats, using a searchlight analysis over an existing fMRI dataset (Tarhan and Konkle, 2020b).

To preview, we found that intuitive action similarity judgments are best predicted by the actors’ goals, followed by the actors’ movement kinematics. Further, our opportunistic analysis of existing fMRI data did not reveal any localized regions with a response similarity structure that was highly correlated with these intuitive similarities. However, we did find tentative evidence for a hierarchical gradient of action processing in the visual system, starting with appearance-based similarity in the early visual cortex, through movement-based similarity in the lateral occipito-temporal cortex, extending to goal-based similarity in the temporo-parietal junction. These data thus highlight an action processing hierarchy within a single, naturalistic action dataset. Overall, these findings suggest that humans are naturally tuned to process others’ intentions, and to a lesser extent their kinematic properties, when observing their actions.

## RESULTS

### Intuitive Action Similarity Judgments

To investigate the principles guiding the perception of a wide variety of actions, we used videos from an existing dataset, depicting 60 everyday actions (Tarhan and Konkle, 2020b). These actions were selected from the American Time Use Survey (U.S. Bureau of Labor Statistics, 2014), which records the activities that Americans typically perform. We chose actions that spanned a range of familiar, everyday activities – such as cooking, running, and laughing – and that engaged objects, people, and their surroundings. Most actions involved a single agent, but a small subset involved two interacting agents (e.g., shaking hands). We selected one short (2.5-second) video to depict each of these actions (see **Methods**). These videos thus depict a wide range of actions, from hand-centric tool actions, like knitting, to aerobic actions that engage the whole body, like dancing. In addition, they are richly varied in their backgrounds, actors, and lower-level motion features such as speed and direction.

We measured intuitive similarities among these videos using a multi-arrangement task adapted from Kriegeskorte and Mur (2012) (**Figure 1a**, see **Methods**). In this task, participants watched all 60 action videos. Then, they saw a blank white circle surrounded by representative still frames from each video. They were told to drag these still frames into the circle, then arrange them according to their similarity: stills from videos that seemed similar were placed closer together, while stills from videos that seemed different were placed further apart (Kriegeskorte and Mur, 2012). We intentionally gave participants very minimal instructions about how to judge similarity; instead, we asked them to do so based on their natural intuitions.

**Figure 1:**
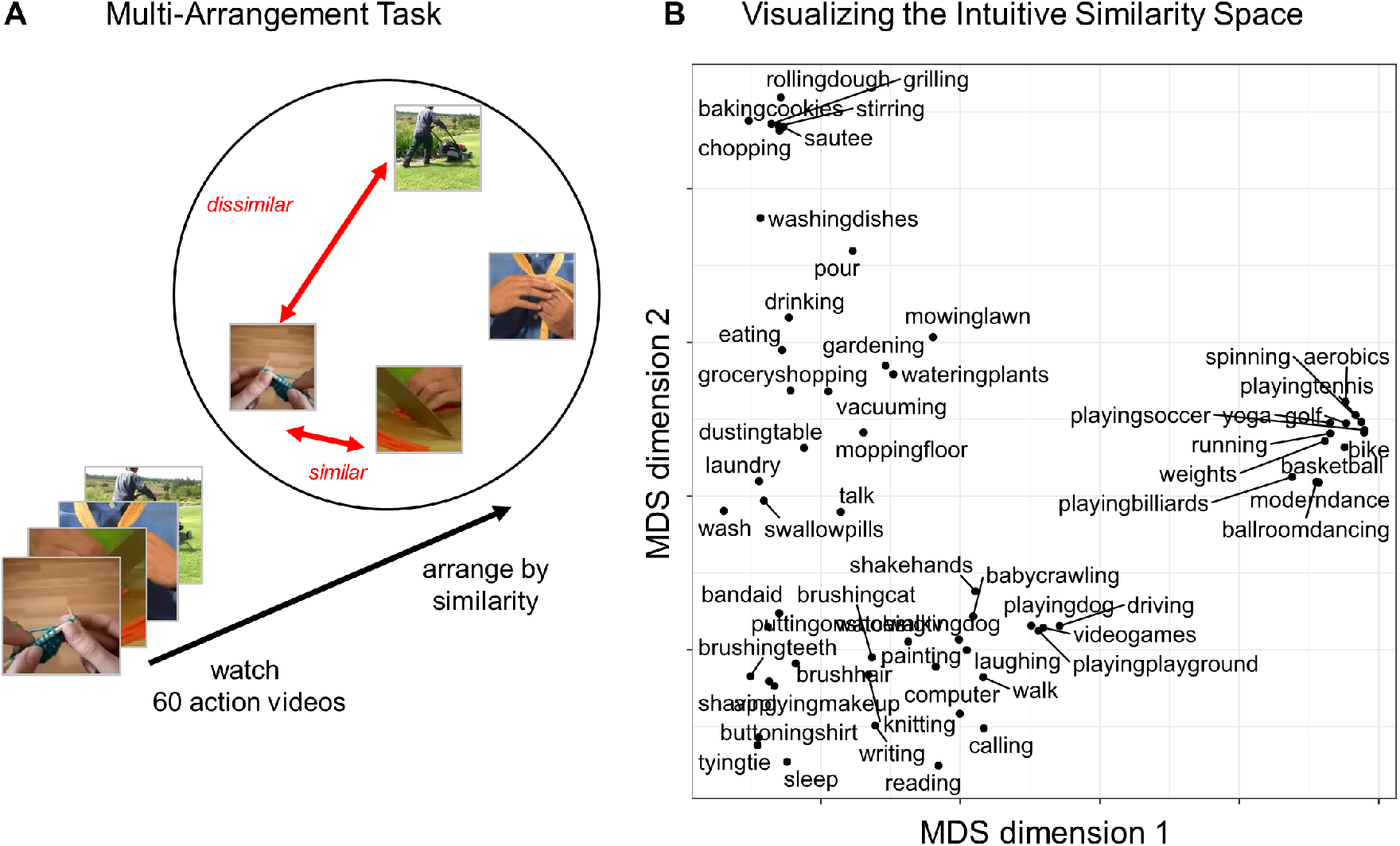
Measuring Intuitive Action Perception. (A) To measure intuitive judgments of action similarity, participants completed an action arrangement task, during which they watched the 60 action videos and then arranged key frames from the videos according to their similarity: frames were close together if participants thought the videos were similar, or far apart if they thought they were different. (B) Plot of the first two dimensions of a Multi-Dimensional Scaling projection, to visualize broad trends in the structure in these intuitive judgments. Actions are plotted close together in this projection if participants consistently judged them to be similar.

We measured these intuitive similarity judgments in one main experiment (Experiment 1; *N* = 19) and in one replication with new participants (Experiment 2; *N* = 20). The group-level similarity judg ments were very consistent across experiments (*r* = 0.89). We also assessed the inter-subject reliability for each experiment by iteratively dividing the participants into two groups and then correlating the averaged data across groups (see **Methods**). In both experiments, we found moderate inter-subject reliability (average split-half Kendall’s τ-a correlation with Spearman-Brown Prophecy correction = 0.51 (Experiment 1), 0.57 (Experiment 2)).

To visualize the structure in these intuitive similarity judgments, we projected the group-level similarities for Experiment 1 into two dimensions using multi-dimensional scaling (MDS). **Figure 1b** shows the resulting projection. The configuration of the actions in this projection provides some insight into what kinds of actions participants regarded as similar. For example, sports actions (e.g., basketball), cooking actions (e.g., chopping), and getting-ready actions (e.g., tying a tie) were all placed in distinct clusters in this projection.

### Predicting Intuitive Similarity with Guided Behavior

To understand the structure of these intuitive action similarity judgments more quantitatively, we used a predictive modeling framework to test what kind of properties could predict perceived action similarity the best. To capture properties at different levels of abstraction, we collected guided similarity judgments based on three different dimensions: the videos’ visual appearance, the actors’ movements, and the actors’ goals. To gather these judgments, we asked new participants to arrange the 60 action videos using the same similarity-based arrangement paradigm as before, but with more explicit instructions. One group of participants (*N* = 20) was told, “please arrange these still images according to their overall visual similarity,” regardless of the actions being performed. This group was encouraged to consider details like the colors in the scene and the direction of the actors’ movements. A second group (*N* = 20) was told to arrange the videos based on similarity in the “actors’ body movements.” This group was encouraged to consider details like whether the actors made large or small movements, moved smoothly or suddenly, and what body parts they were using. Finally, a third group (*N* = 20) was told to arrange the videos based on similarity in the “actors’ goals.” These tasks were meant to capture relatively lower-level visual properties, more intermediate-level kinematic properties, and higher-level inferences about the actions. The group-level representational dissimilarity matrices (RDMs) from these three experiments operationalized our three hypotheses for the kinds of information that underlie intuitive action similarity judgments. **Supplemental Figure 1** displays the resulting RDMs and their correlations. Hereafter, we refer to these matrices as “model RDMs.”

We then asked how well each model RDM predicted the intuitive similarity judgments. To do this, we used linear regression: each model RDM was entered into a separate regression to predict the intuitive similarity judgments. To estimate the best possible prediction performance, we calculated the data’s noise ceiling as a range between the 25th and 75th percentiles of the data’s split-half reliability (Spearman-Brown Prophecy-corrected Kendall’s τ-a = 0.50 – 0.55 (Experiment 1), 0.56-0.59 (Experiment 2)). We assessed prediction performance for each model RDM using a leave-1-action-out cross-validation procedure: the regression was iteratively trained on all intuitive similarity judgments except those involving one held-out action (e.g., the 1,711 similarities between action pairs not involving running), then tested on the held-out data (the 59 similarities between running and all other actions). Prediction accuracy was calculated by correlating the actual and predicted intuitive similarities for this held-out data. If a model had high prediction accuracy, then the properties that it captures might underlie intuitive action similarity judgments.

**Figure 2a** and **Table 1** show the results of these analyses. In general, the model RDM based on the actors’ goals predicted the data very well, while the model RDMs based on the actors’ movements and the videos’ visual appearance both predicted the data moderately well (**Figure 2a**). This conclusion was supported by a 2 × 3 (experiments x model RDMs) ANOVA, which revealed a significant main effect of model RDM (F(2, 354) = 45.9, *p* < 0.001). Post-hoc comparisons indicate that the visual and movement model RDMs did not differ, but both performed significantly worse than the goal model RDM (**Table 1**). There was no main effect of experiment (F(1, 354) = 0.21, *p* = 0.65) and no interaction (F(2, 354) = 0.41, *p* = 0.66). Altogether, these results show that the actors’ goals predicted intuitive similarity judgments the best of the three hypothesized properties, but the actions’ movements and the videos’ visual appearance also predicted these intuitive judgments relatively well.

**Table 1:**
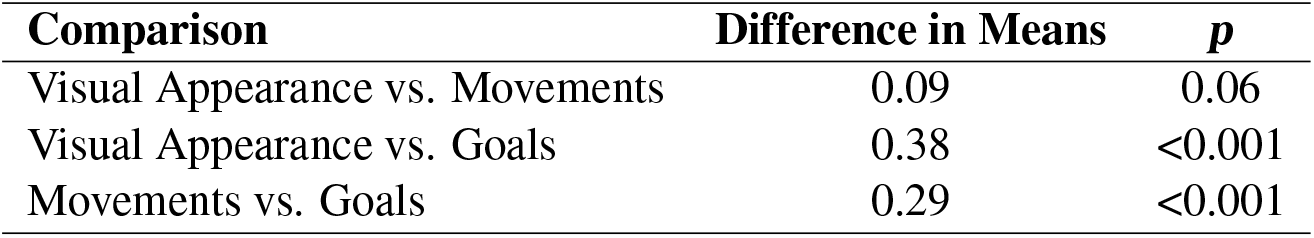
Results of post-hoc tests investigating the main effect of model RDM.

**Figure 2:**
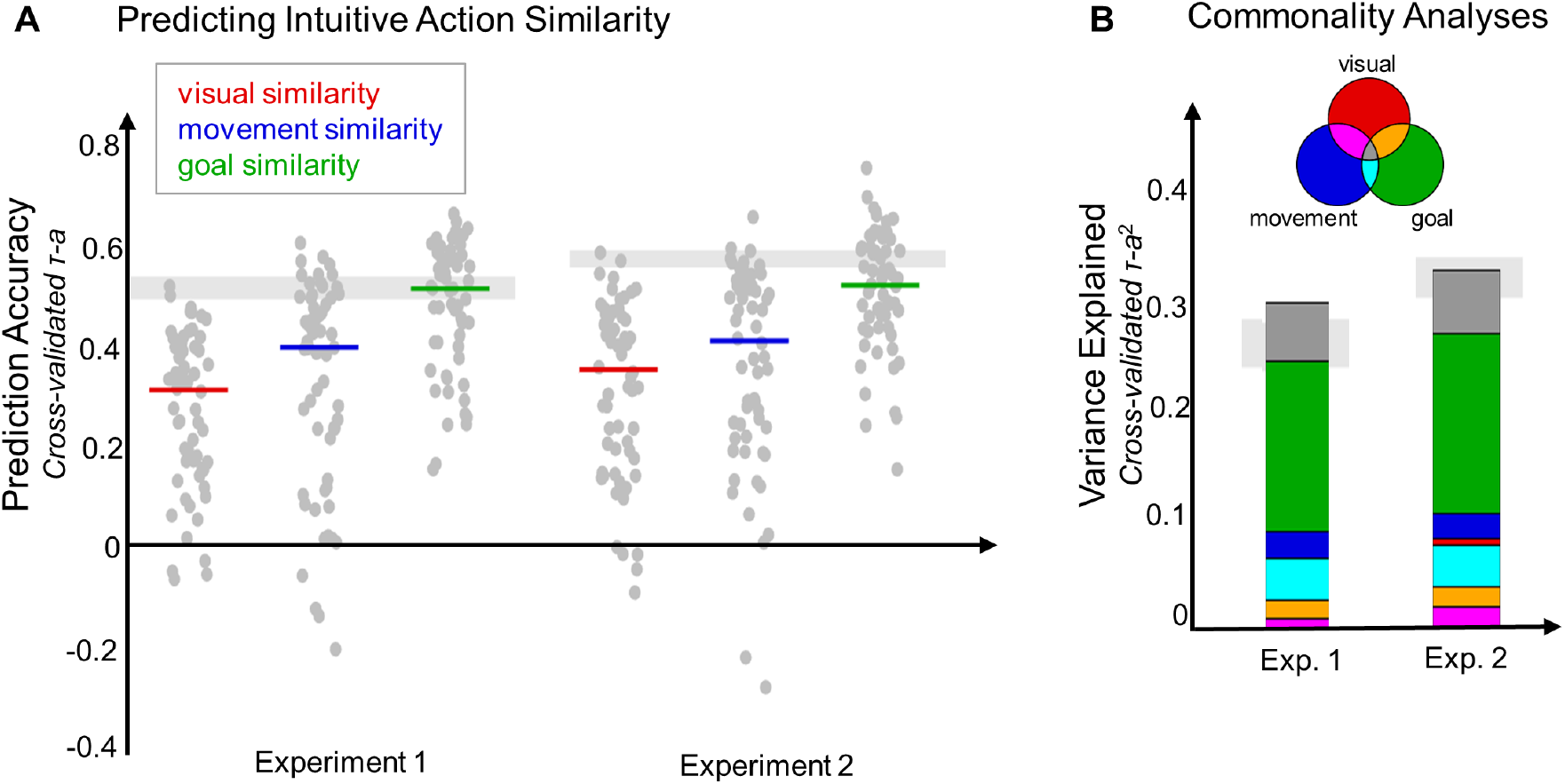
Predicting Intuitive Action Similarities. (A) To investigate what kinds of properties underlie intuitive action similarities, we tested how well similarity along three broad dimensions (model RDMs) predicted the intuitive judgments. Prediction performance was measured for each model RDM in a leave-1-condition-out cross-validation procedure. The results are shown for each experiment. Light grey bars indicate the noise ceiling for each experiment. Colored horizontal lines indicate the median prediction accuracy for each model RDM, calculated over all iterations of the leave-1-condition-out procedure. Grey dots show the prediction performance on each iteration of this procedure (1 dot per held-out condition). (B) Commonality analyses (Lescroart et al., 2015) were used to assess how much of the variance explained in the intuitive similarity judgments was shared between the model RDMs, and how much was unique to one model RDM. Venn diagram illustrates the meaning of each color: for example, variance that the goal similarity judgments uniquely explained is shown in green, variance shared between the goal and movement similarity judgments is shown in light blue, and variance shared by goal, movement, and visual similarity judgments is shown in dark grey. Light grey bars indicate the maximum explainable variance (noise ceiling^2^) for each experiment.

These results raise a natural question: how unique are these three model RDMsã Do the similarity judgments based on visual appearance and movements account for different components of the intuitive similarity judgments, or do they overlapã To address this question, we performed a commonality analysis (Lescroart et al., 2015) to assess how much variance in the intuitive similarity judgments was uniquely accounted for by each model RDM, and how much was shared between model RDMs. This analysis was particularly crucial to understanding what kinds of information influence the intuitive similarity judgments, because the model RDMs captured some overlapping information (**Supplemental Figure 1**).

**Figure 2b** and **Table 2** show the results of this analysis. In general, the actors’ goals accounted for the most unique variance (green bars in **Figure 2b**), indicating that this information is sufficient to explain a large portion of the explainable variance in the intuitive similarity judgments. The actor’s movements accounted for a much smaller amount of unique variance, while the videos’ visual appearance did not account for any unique variance (see **Table 2** for the complete set of results). In combination, these three model RDMs accounted for roughly all of the explainable variance in the intuitive similarity judgments (Experiment 1: 31% / 24%; Experiment 2: 34% / 31%). Note that these models explained slightly more than the maximum explainable variance, which is possible because this is only an estimate of the true maximum. Altogether, these results suggest that the actors’ goals not only predicted the intuitive similarity judgments the best of the three model RDMs that we tested; they also accounted for far more unique variance in the data.

**Table 2:** Commonality Analyses. Results of commonality analyses to investigate how much variance in the intuitive similarity judgements each model RDM uniquely accounts for, and how much is shared between model RDMs. The maximum explainable variance (noise ceiling^2^) was 24% for the main experiment and 31% for the replication.

| Experiment | Partition of the Variance | % Explained |
| --- | --- | --- |
| Main Experiment | Unique to Visual Appearance | -0.02 |
|  | Unique to Movements | 2.6 |
|  | Unique to Goals | 16.0 |
|  | Visual Appearance & Movements | 1.0 |
|  | Visual Appearance & Goals | 1.7 |
|  | Movements & Goals | 4.0 |
|  | Visual Appearance & Movements & Goals | 5.5 |
| Replication | Unique to Visual Appearance | 0.6 |
|  | Unique to Movements | 2.4 |
|  | Unique to Goals | 17.0 |
|  | Visual Appearance & Movements | 1.9 |
|  | Visual Appearance & Goals | 2.0 |
|  | Movements & Goals | 3.9 |
|  | Visual Appearance & Movements & Goals | 5.9 |

### Neural Correlates of Intuitive Action Similarity

In a previous study, we collected functional neuroimaging data while a separate set of participants viewed these same videos (Tarhan and Konkle, 2020b). Here we take advantage of this existing dataset to conduct opportunistic exploratory analyses to examine if there are any regions that show strong correspondence with the intuitive measure of action similarity. Note that in this paradigm, participants passively viewed the videos–no explicit similarity relationships among videos were task-relevant in this experiment.

To assess where in the brain, if anywhere, there is a match between the intuitive action similarity judgments and localized neural similarity structure, we conducted a whole-brain searchlight representational similarity analysis (RSA; Kriegeskorte et al., 2006). This analysis compared the intuitive judgments to representational dissimilarities within circumscribed searchlight spheres centered at each voxel in the cortex.

As our first analysis step, we estimated and visualized how reliable the data in these brain search-lights were. To do so, we calculated each searchlight’s split-half reliability, correlating the neural RDMs from odd-numbered imaging runs with RDMs from even-numbered runs (see **Methods**; Tarhan and Konkle, 2020a). The resulting reliability map (**Figure 3a**) shows that the searchlight data are quite reliable in the dorsal and ventral streams of the visual cortex (max. split-half *r* = 0.88); however, reliability is low in the prefrontal cortex, anterior temporal lobe, and medial parietal cortex (min. split-half *r* = -0.13). This reliability map serves as a useful guide for interpreting the RSA results: the low reliability outside of the visual cortex meant that we could not expect to find strong brain-behavior correlations in those regions. However, the data had enough signal to observe strong correlations in much of the visual cortex.

**Figure 3:**
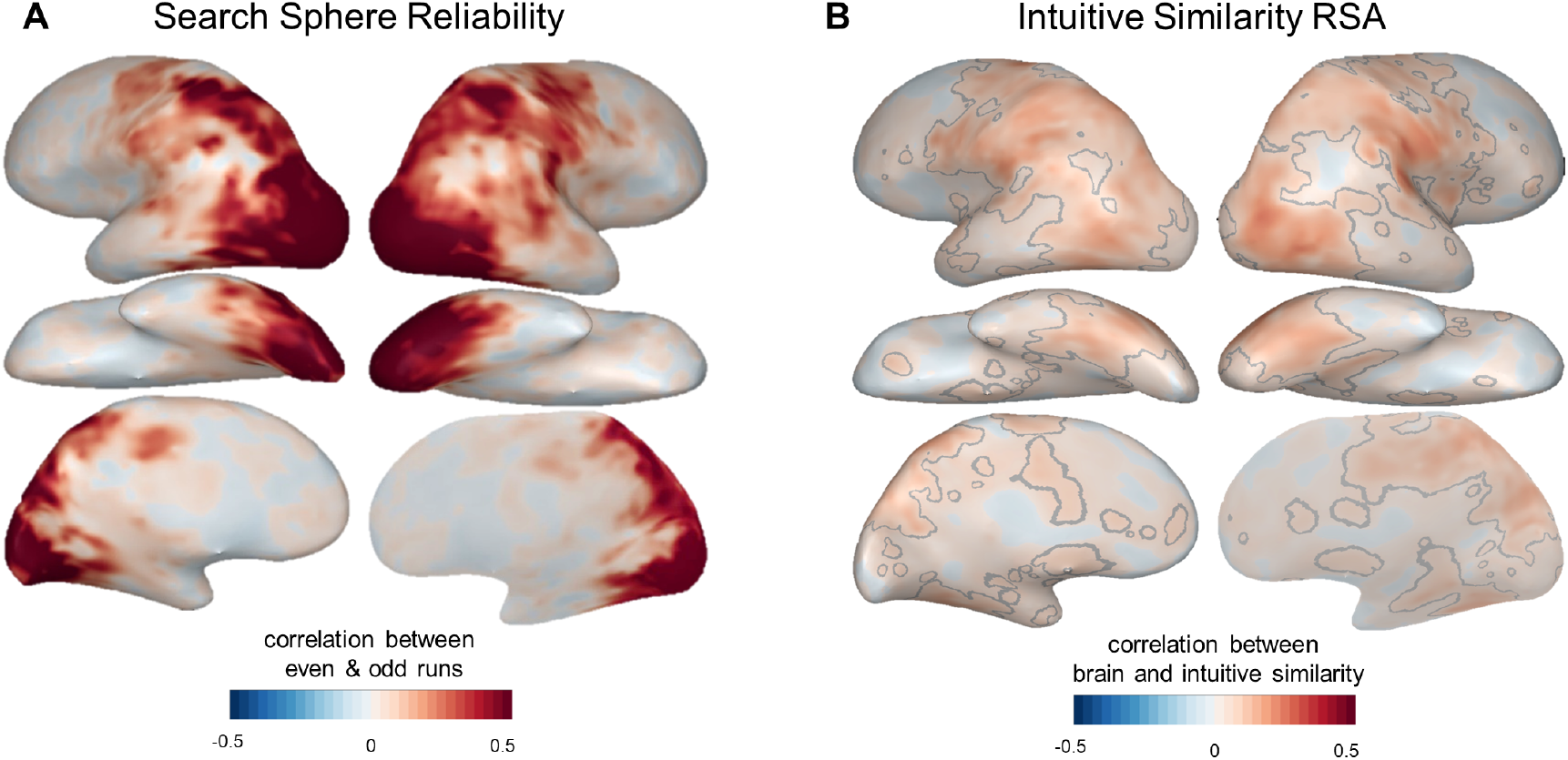
Exploratory Searchlight Analysis. Whole-brain searchlight analyses were used to compare intuitive similarity judgments to neural responses measured in a separate fMRI experiment (Tarhan and Konkle, 2020b). (A) Searchlight split-half reliability map, showing the correlations between neural dissimilarity matrices calculated based on even- and odd-numbered imaging runs (see Methods). We have higher confidence in the Representational Similarity Analysis results in voxels with higher searchlight reliability. (B) Searchlight results comparing neural response geometries to intuitive judgments of action similarity. Grey lines outline the voxels that survived statistical corrections (voxelwise permutation tests at p < 0.01, cluster-level permutation tests at q < 0.05).

Next, we correlated these searchlight RDMs with the intuitive similarity judgments to determine how well the representational structure in each searchlight captured the intuitive-level structure. As shown in **Figure 3b**, we found significant brain-behavior correlations throughout the ventral and dorsal visual streams, as well as primary somatosensory strip, primary motor strip, premotor cortex, and the medial parietal lobe (the areas outlined in grey, which survived voxel-wise permutations (*p* < 0.01) and cluster-level permutations (*q* < 0.05)). These correlations were strongest along the lateral temporal cortex and superior temporal cortex, in the vicinity of regions that may represent objects’ functions and kinematics (Bracci et al., 2012; Bracci and Peelen, 2013; Leshinskaya and Caramazza, 2015). However, these correlations were relatively weak (mean *r* among significant voxels = 0.11, s.d. = 0.04, range = 0.04-0.30), especially considering that the high reliability in these regions suggested that it should be feasible to find stronger correlations if they exist (maximum split-half reliability *r* = 0.88). Therefore, these results suggest that intuitive similarity judgments do not strongly draw on representations in the visual cortex; however, they also leave open the possibility that these judgments draw more strongly on representations in the prefrontal or anterior temporal cortex. We discuss this and other possibilities in the **Discussion**.

We next examined how well our three hypothesized model RDMs could account for these response similarities. Specifically, we examined whether the relatively low, intermediate, and higher-level action properties would account for the neural response structure in increasingly high-level regions of the visuo-motor cortex, which would reveal a hierarchical gradient of action-related processing starting in the visual system.

To investigate this possibility, we first conducted whole-brain searchlight RSAs to compare each model RDM to the brain (**Figure 4**, left panel), with significant relationships outlined in grey (voxel-wise permutation tests: *p* < 0.01 with permutation-based cluster corrections *q* < 0.05). These model-searchlight results reveal that much of the ventral and dorsal stream has local regional similarity structure that corresponds with the three different similarity measures, to different degrees.

**Figure 4:**
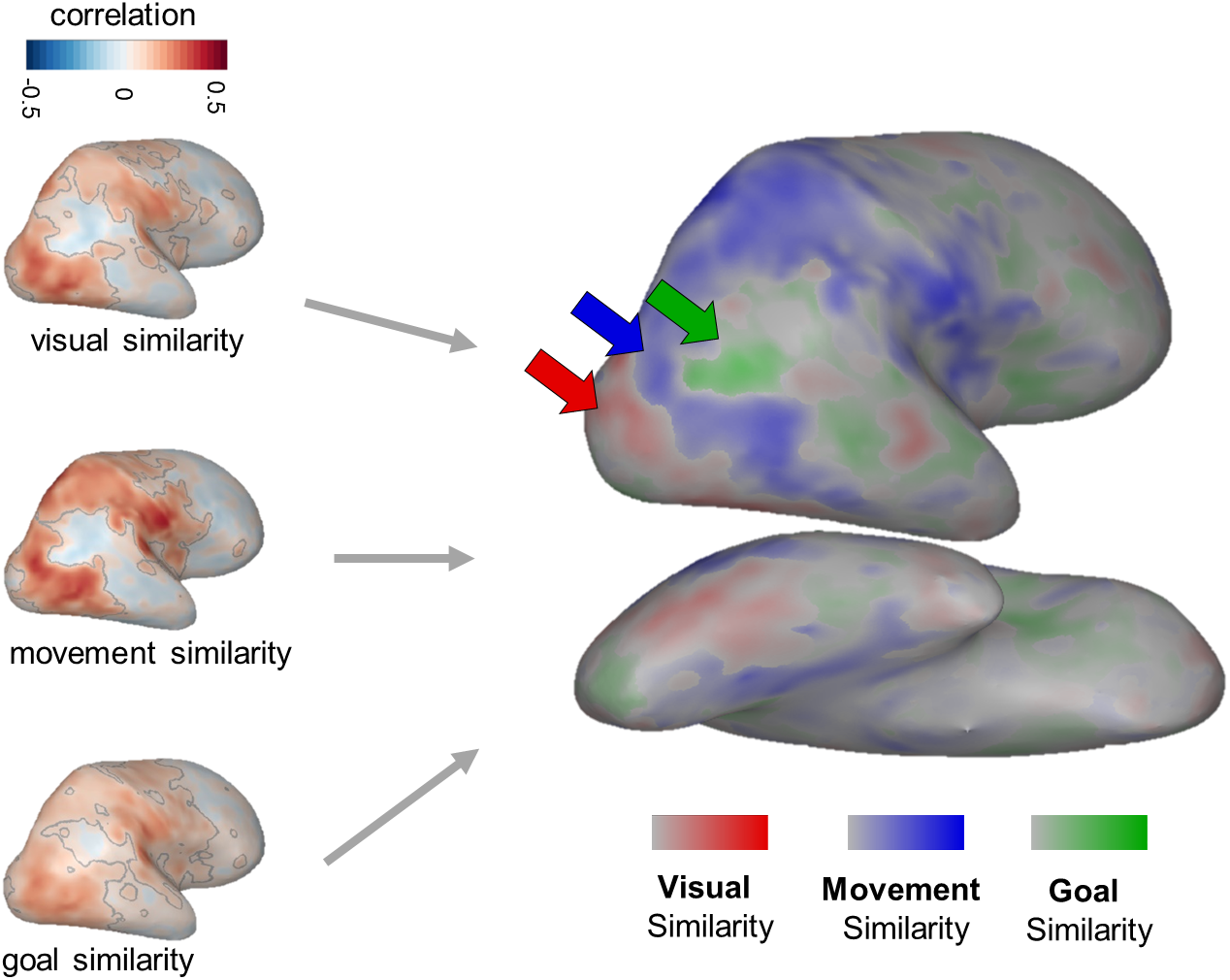
Visualizing the Action Processing Hierarchy. To understand which brain regions are most related to each of the hypothesized kinds of action similarity, separate Representational Similarity Analyses were conducted comparing each of the three model RDMs to neural responses. RSA results for each model RDM are shown on the left, with significant voxels outlined in grey. A 3-way winner map (right) was calculated by identifying the model RDM with the highest positive correlation to each searchlight and coloring the searchlight’s central voxel according to that RDM. The intensity of the color indicates the difference between the strongest and next-strongest correlation.

To visually compare the general topographic distribution of the strength of these searchlight correlations over the entire brain, we calculated a 3-way winner map, in which each voxel is colored according to the model RDM that was most strongly correlated with the searchlight RDM centered at that voxel. Only voxels with positive model-brain correlations are plotted. Note that no statistical tests were conducted over the topographic distribution or strengths of this 3-way winner map visualization. Thus these results should be interpreted cautiously, and are well-suited for deriving more specific hypotheses about the cortical locations with different representational formats, that require further testing for confirmation.

This winner map shows some suggestive evidence for a hierarchical progression in the structure of brain responses. That is, the representational structure in the early visual cortex and parts of right ventral temporal cortex is best captured by the model RDM based on visual appearance (red), while the structure in the lateral temporal cortex and intra-parietal sulcus is best captured by the model RDM based on movement kinematics (blue). Finally, regions known to be involved in social processing, including the right temporo-parietal junction (TPJ), are captured best by the model RDM based on the actors’ goals (green; Dodell-Feder et al., 2011; Koster-Hale et al., 2017; Pitcher and Ungerleider, 2020; Saxe et al., 2004). Note that the TPJ lies just outside of our reliable coverage (**Figure 3a**), so while this result aligns with strong prior evidence in the literature, we avoid drawing strong conclusions from this analysis about the representations in this region.

The progression of best matching models highlights that actions are represented based on different properties at different stages of visual processing. While this hierarchy echoes previous work on the structure of the visual system, to our knowledge this is the first time it has been shown (i) specifically for action processing, and (ii) in a single, naturalistic action dataset. It is also notable that some model RDMs were more strongly correlated with the representational structure in the visual cortex than the intuitive similarity judgments were (e.g., max. *r* = 0.45 for movement similarity judgments, compared to 0.3 for intuitive judgments). This further suggests that there was enough signal in the data from these regions to pick up on a stronger correlation between the brain and the intuitive similarity judgments, if one existed.

Altogether, these analyses suggest that the representations underlying intuitive action similarity are not cleanly localized to a circumscribed region within action-responsive visuo-motor cortex. Rather, we found suggestive evidence for an action processing hierarchy that unfolds across the visual cortex and likely extends into social-processing regions in the temporo-parietal junction.

## DISCUSSION

Here we investigated how well properties at different levels of abstraction capture behavioral judgments about the intuitive similarities among a large set of everyday action videos. We found that the actors’ goals strongly predicted these intuitive similarities, while the actors’ movements also contributed to these judgments, but visual appearance contributed little to nothing. These findings add to existing evidence that humans naturally process others’ motivations when they observe and compare their actions. To add to this cognitive investigation, we found evidence for a representational gradient in the brain, whereby early visual cortex represents actions’ visual appearance and higher-level visual cortex represents more intermediate-level kinematic information. This gradient highlights transitions in the structure of action representations along the visual processing stream; notably, we found this representational gradient in a single, naturalistic dataset. In the following sections, we situate these findings in the literature, highlight how this work advances existing methods for understanding action processing, and discuss promising next steps.

### Intuitive Action Representations in the Mind

Our primary finding was that judgments about the similarity of actors’ goals was the best predictor of intuitive action similarity judgments. In addition, these goals accounted for the most unique variance in the intuitive similarity data. We interpret this to mean that humans naturally and intuitively process other actors’ internal motivations and thoughts, even in the absence of an explicitly social task. This conclusion adds to a rich literature showing that humans automatically represent others in terms of their mental states, even from a very young age (Gergely and Csibra, 2003; Jara-Ettinger et al., 2016; Liu et al., 2017; Reid et al., 2007; Thornton et al., 2019a,b). In addition, we found that similarity in the actors’ movements also predicted intuitive judgments moderately well and accounted for a smaller amount of unique variance in the data. This finding goes beyond our current understanding of the factors driving natural action processing, to suggest that kinematic information also contributes to intuitive action perception. In contrast, similarity in the videos’ visual appearance did not account for any unique variance in the data, suggesting that lower-level visual properties such as color, form, and motion direction do not have much influence on natural action perception.

It is important to note that the multi-arrangement task used here can tell us which actions are more similar than others based on a instructed property, but not *why* they are similar. Thus, a natural extension of these findings is to investigate the format of these goal- and movement-based representations–that is, what specific features do humans consider when estimating the similarity in actors’ goals and movements? For example, how important are speed, trajectory, and movement quality (e.g., shaky or smooth) for our assessment of the similarity among actions’ movements? Do we consider physical variables – such as facial expression – when inferring actors’ goals? Recent empirical advancements provide concrete methods for addressing these questions. For example, modeling approaches that learn sparse feature-based representations allow researchers to infer the format and dimensionality of the representations underlying similarity judgments (Hebart et al., 2020). Additionally, advances in deep learning models provide tools to explore whether image- and video-computable feature spaces match behaviorally-measured similarity spaces.

Related to this point, we used behavioral judgments, rather than image-computable features, to estimate a similarity based on visual appearance. But, one concern is whether behavioral similarity judgments can even capture low-level visual properties—do individuals have explicit access to this level of representation? Further, if individuals spontaneously represent actions in terms of their goals, how can we be sure that the similarity judgements based on visual and movement properties were not affected by the spontaneous processing of the actors’ goals? These are important questions because the logic of our design assumes that people can flexibly and accurately report different kinds of similarity relationships, given explicit instructions. Empirically, the fact that the three kinds of similarity judgments showed distinct and reliable variance (**Supplementary Figure 1**) supports the validity of this assumption. Additionally, the fact that these behaviorally-measured similarity spaces showed some correspondence with brain similarity structure (e.g. with visual appearance similarity showed the strongest correspondence with the early visual cortical regions) further supports this logic. In general, a strength of our empirical approach is that using the same behavioral task with different targeted instructions allows these model similarity spaces to be more comparable in their format, and the similarity spaces are clearly also behaviorally-relevant.

### Intuitive Action Representations in the Brain

In addition, we explored the neural basis of intuitive action perception. In an opportunistic representational similarity analysis, we found that the representational geometries in regions in the lateral occipito-temporal, intra-parietal, and sensorimotor cortices were only weakly correlated with intuitive similarity judgments—we did not find strong evidence that these regions support intuitive action representations in a localized manner. Why? One possibility is that intuitive similarity judgments rely on representations that are distributed among a widespread network of brain regions, which would only be detected by a larger-scale analysis – such as decoding from a much larger swathe of cortex. Another possibility is that these results were influenced by the task done in the fMRI scanner. In our data, observers passively viewed the videos. Perhaps a different task – such as making intuitive similarity judgments between successive videos, for example – would reliably engage additional regions with a more goal-based similarity structure, or even modulate the similarity structure measured in the visuo-motor cortex.

Keeping these caveats in mind, in the current neuroimaging dataset there seems to be a division between intuitive action judgements, which relies on fairly abstract information about the actor’s mental states and goals, and the visuo-motor cortex, which represents a range of action properties that may be the perceptual precursors to higher-level processing. Consistently, earlier work by Lestou et al. (2008) also found that areas in the visual and parietal cortex were relatively more sensitive to the kinematics of actions, than to their goals. And more recently, Pitcher and Ungerleider (2020) have proposed that the visual cortex contains a major processing stream dedicated to processing others’ movements. This stream sits between the classic “what” and “where” pathways (Mishkin et al., 1983), and is thought to process the visual information that eventually feeds into more abstract action representations outside of the visual cortex.

This hypothesis suggests that relatively perceptual properties may explain the structure in the visuo-motor cortex well, but fall short of predicting intuitive judgments well. In line with this idea, we found that the visual cortex was more strongly correlated with actions’ visual appearance and movements than it was with the intuitive similarities (**Figure 3b** & **Figure 4**). In addition, our previous work (Tarhan and Konkle, 2020b) indicated that the body parts and targets involved in an action, whether an action is directed at a person (*sociality*), and the scale of space at which it affects the surroundings (*interaction envelope*) all predict responses in this cortex well. In contrast to their prominence in the visuo-motor cortex, these properties only predicted intuitive similarity judgments moderately well (**Supplemental Figure 2**). The perceptual precursors computed en route to intuitive action representations may also include some functional information (such as how bodies and object interact, or whether the action creates something new), which explains action responses in the parietal and lateral occipito-temporal cortices (Thornton and Tamir, 2019; Bracci et al., 2012; Bracci and Peelen, 2013; Leshinskaya and Caramazza, 2015; Tucciarelli et al., 2019). While this functional information may seem more abstract than an action’s visual appearance, it may still be less abstract than the mental state information that scaffolds intuitive similarity judgments. Altogether, this evidence supports the notion that the visuo-motor cortex computes the perceptual precursors of the higher-level representations over which intuitive action perception operates.

Where, then, does the brain house these intuitive action representations? Given that the actors’ goals predicted intuitive judgments very well, it is likely that the answer lies in regions involved in representing others’ mental states or personal attributes. These include the medial prefrontal cortex (mPFC), the anterior temporal lobe (ATL), and the temporo-parietal junction (TPJ; Dodell-Feder et al., 2011; Koster-Hale et al., 2017; Samson et al., 2004; Saxe et al., 2006; Thornton and Tamir, 2019). Others have implicated regions in the ventral premotor cortex (e.g. Lestou et al., 2008, see also Sitnikova et al., 2014; Lingnau and Petris, 2013). We did not find strong correlations between any of these regions and the intuitive similarity judgments. However, this does rule out the possibility that these regions are involved in intuitive action perception. Recall that our reliability analysis revealed that our data are very reliable in the visuo-motor cortex, but much less reliable in the mPFC, ATL, and TPJ. This pattern of reliability makes it virtually impossible to find strong correlations in these social-processing regions, even if they truly are involved in intuitive action perception.

Interpreting our results in light of reliability sets this work apart, because it allows us to qualify which results are informative, and which are not. In any fMRI study, some regions will be more reliable than others (Tarhan and Konkle, 2020a; Eklund et al., 2016). In our case, if we had not accounted for these variations in reliability, we might have concluded that parts of the visual cortex are moderately related to intuitive action perception, while mPFC and ATL are not related at all. In contrast, when we account for these variations, we conclude that these parts of visual cortex are most likely *not* related to intuitive action perception, while mPFC and ATL *may* be related to these intuitions. This is because reliability in the visual cortex regions was so high that we could have found much stronger correlations if they existed. In contrast, reliability in mPFC and ATL was too low to pick up on correlations even if they existed. This difference in our interpretations before and after taking reliability into account highlights the importance and power of reliability analyses for interpreting cognitive neuroscience results.

## METHODS

### Data Availability

All data, stimuli, and main analysis scripts are available on the Open Science Framework repository for this paper (https://osf.io/d5j3h/).

### Experimental Procedures

#### Participants

113 participants were recruited through the Harvard Psychology Department for in-lab action arrangement studies. These participants were either paid $15 or given course credit. Data were excluded from 15 participants because of incomplete or unreliable data (participant-level noise ceilings < 0.1). All participants gave informed consent in accordance with the Harvard University Institutional Review Board and the Declaration of Helsinki.

#### Stimuli

The stimuli consisted of 60 videos depicting everyday actions (Tarhan and Konkle, 2020b). The 60 actions were selected from the American Time Use Survey (U.S. Bureau of Labor Statistics, 2014), which surveyed a large sample of Americans about the activities that make up their days. The videos were edited to 2.5 seconds in duration, with a square (512 × 512 pixel) frame centered on the main actor.

#### Multi-Arrangement Task

In two experiments, participants (Experiment 1: *N* = 19, 7 males, mean age: 21.5 years; Experiment 2: *N* = 20, 6 males, mean age: 21.7 years) completed a multi-arrangement task adapted from Kriegeskorte and Mur (2012) (**Figure 1a**). First, they watched all 60 videos without sound. The videos were played in a randomized order without breaks in full-screen mode (monitor dimensions: 18.75 × 10.5 inches). Once the videos had finished playing, the multi-arrangement task began. On each trial of this task, participants saw a blank white circle surrounded by key frame images from the videos. They were told to drag the images into the circle, then arrange them so that images from similar videos were closer together and images from different videos were further apart. They were also told that there was no “right” way to arrange the videos; rather, they should use their intuitions to decide how similar the videos were. Once they had arranged all of the key frame images in the circle, they could continue on to the next trial; there was no time limit for each trial. In addition, they could re-play any video as much as they wanted in a separate window, to remind themselves of what it looked like.

In order to collect reliable data in an efficient way, we used a “lift-the-weakest” algorithm (Kriegeskorte and Mur, 2012) to determine which key frames to show on each trial. On the first trial, participants arranged key frames from all 60 videos. Then, they completed approximately 20-70 subsequent trials where they arranged key frames from a sub-set of the 60 videos. The algorithm selected key frames just before each trial, based on an accumulated evidence criterion (signal-to-noise ratio^2^). Evidence scores were calculated for each pair of videos after each trial. Pairs received a low evidence score if the actions had not been arranged relative to each other many times, or if they had been arranged inconsistently during prior trials. Key frames from these actions were more likely to be presented in subsequent trials, in order to measure their perceived similarities more accurately. Often, low evidence action pairs had been placed very close to each other within a cluster – focusing on these clusters during subsequent trials allowed us to capture the finer-grained distances among actions within a cluster. The trials continued until all action pairs achieved a minimum evidence criterion of 0.5 or the experiment timed out (after 60 minutes, excluding time for breaks).

At the end of the experiment, the data consisted of the lower triangle of a distance matrix between all action videos. Each cell (*i, j*) contained the estimated Euclidean distance between videos *i* and *j*, built up over trials. This estimate was calculated using an inverse multi-dimensional scaling algorithm to infer distances between videos that were presented on-screen in different sub-sets. In addition, these distances were normalized to account for the fact that different trials presented different numbers of key frames within the same amount of screen space. More details on the lift-the-weakest algorithm, inverse multi-dimensional scaling, and normalizing procedures can be found in Kriegeskorte and Mur (2012).

#### Guided Similarity Judgments

To measure the actions’ similarity according to specific kinds of properties, we also ran a variant of the multi-arrangement task with more explicit instructions, with new participants. One group (*N* = 20) was instructed to arrange key frame images from the 60 action videos according to similarity in their overall visual appearance. Specifically, they were told, “Please arrange these still images according to their overall visual similarity, regardless of the actions in the videos”, and during the practice trials the experimenter encouraged participants to take information like the colors and the direction of the movements into account when arranging the actions. Another group (*N* = 20) was instructed to arrange the key frames according to similarity in the actors’ manner of movement. This group was told, “Please arrange these still images according to the actors’ body movements”, and during the practice trials the experimenter encouraged them to pay attention to the body parts being used, the amount of movement in the video, whether it was smooth or abrupt, et cetera. A third group (*N* = 20) was instructed to arrange the key frames according to similarity in the actors’ goals. They were told, “Please arrange these still images according to similarity in the actors’ goals.” Distance matrices were averaged across participants for each of these conditions, producing three model RDMs that were used to predict the intuitive similarity judgments measured in the multi-arrangement task.

#### fMRI Data

We used data from Tarhan and Konkle (2020b) to analyze neural responses to the same action videos as were used in the multi-arrangement task. In that experiment, 13 participants completed a 2-hour fMRI scanning session, during which they passively viewed the videos and detected an occasional red frame around the videos to maintain alertness. Further details about these data can be found in Tarhan and Konkle (2020b) and at the paper’s Open Science Framework repository (https://osf.io/uvbg7/)

#### Multi-Dimensional Scaling Analysis

Multi-Dimensional Scaling (MDS) was performed over the intuitive similarity judgments from Experiment 1, to visualize the overall structure in these judgments. The distance matrices measured in the multi-arrangement task were averaged across individual participants and non-metric MDS was performed over this group-averaged distance matrix in MATLAB. We extracted the first two dimensions of the resulting projection and plotted them as a scatterplot (**Figure 1b**). Note that we extracted two dimensions for ease of visualization, but stress plots indicated that four dimensions would more fully capture the structure of the data.

### Modeling Analyses

#### Noise Ceilings

We used a split-half procedure to calculate the noise ceiling for the intuitive similarity judgements, to provide a reference for how well we could expect any model to predict the intuitive similarity judgments given the data’s inherent noise. We randomly divided individual participants into two groups, then averaged the distance judgments over all participants in each group and calculated the Kendall’s τ-a correlation between the groups. This procedure was repeated 100 times, to build up a distribution of split-half correlation values. We then corrected for the effects of splitting the data by applying a Spearman-Brown Prophecy Correction. We estimated the noise ceiling as a range from this distribution’s 25^th^ percentile to its 75^th^ percentile.

#### Predictive Modeling

We used cross-validated regression to assess how well the three model RDMs – the judgments about the videos’ visual appearance, movements, and the actors’ goals – could predict the intuitive similarity judgments. First, we averaged the intuitive similarity judgments across all participants, and both these and the model RDMs were z-normalized so that they had a mean value of 0 and a standard deviation of 1. Then, we iteratively fit an Ordinary Least Squares regression to the intuitive similarity judgments for each model RDM. On each iteration, we held out the data from one action (distances between 59 pairs of actions) and fit the model using the data from 59/60 videos (1,711 pairs). We then calculated the predicted intuitive similarity judgments for the held-out data using the weights fit on the training data. Because each pair of actions was held out twice during this cross-validation procedure (once when holding out all pairs involving action i and again when holding out all pairs involving action j), we averaged over the two predicted intuitive similarities to obtain a single predicted intuitive similarity judgment for each pair. Finally, to assess how well each model RDM predicted the held-out data across these iterations, we correlated the predicted similarity judgments with the actual similarity judgments using Kendall’s τ-a correlation. This correlation was calculated separately for each iteration —-over the pairs that were held out during that iteration —-and then averaged over the iterations. This entire procedure was performed separately for each model RDM and each of the two experiments.

#### Comparing Predictive Models

To compare the model RDMs’ prediction performance across experiments, we conducted a 3×2 (model RDMs x experiments) ANOVA. We accounted for the fact that noise ceilings differed across experiments by re-scaling the prediction results as a proportion of the noise ceiling’s lower bound. Post-hoc tests were run to investigate any significant main effects, using the Tukey-Kramer correction for multiple comparisons.

#### Commonality Analyses

Commonality Analyses were used to assess how much each model RDM’s prediction performance reflected its ability to account for unique variance in the intuitive similarity judgments, and how much was shared with other model RDMs. To do this, we followed the procedure described in Lescroart et al. (2015).

First, we ran seven regressions, using all possible combinations of the three model RDMs to predict the intuitive similarity judgments. That is, we ran one regression predicting the intuitive judgments with all three model RDMs: one with the goal- and movement-based RDMs; one with just the goal-based RDM, and so on. For each regression, we estimated the squared leave-1-condition-out prediction value (Kendall’s τ-a^2^), which is an approximation of the amount of variance explained in the intuitive judgments by the predictors entered into that regression.

To calculate the amount of variance in the intuitive similarities that was uniquely explained by a model RDM (e.g., goal-based similarity), we subtracted the τ-a^2^ value for the combination of the other two model RDMs from the τ-a^2^ value for the combination of all three RDMs. For example, the unique variance (*UV*) explained by goal-based similarity was calculated as:

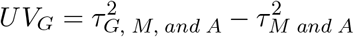

Where

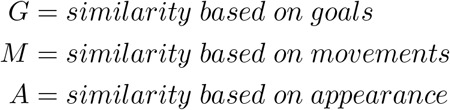

We calculated the amount of variance shared by all three model RDMs (*SV*) as:

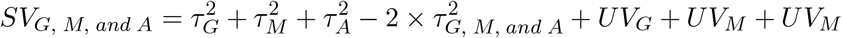

Finally, the amount of variance shared between the goal and movement model RDMs (but not the appearance RDM) was:

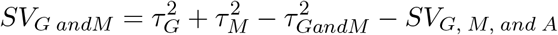

To calculate the total explainable variance in the data, we simply squared the noise ceiling range for each experiment.

#### Whole-Brain Searchlight Analyses

Whole-brain searchlight representational similarity analyses were conducted to map out where, if anywhere, the brain’s representational geometry matches the structure in the intuitive similarity judgments. For each gray-matter voxel, we calculated a neural Representational Dissimilarity Matrix (RDM) based on the responses from gray-matter voxels within 9 mm (3 voxels) of that voxel. On average, each searchlight contained 121.3 voxels (s.d. = 4.5). Neural RDMs were calculated over the voxels in the searchlight using the correlation distance between the response patterns for each pair of actions.

#### Searchlight Reliability

Before comparing these neural RDMs to the behavior, we assessed the reliability of the RDM in each search sphere. To do this, we calculated separate neural RDMs using data from odd- and even-numbered imaging runs. Then, we correlated these splits of the data, resulting in a map of search sphere reliability across the cortex (**Figure 3a**). This procedure is a variation on the one described in Tarhan and Konkle (2020a).

#### Searchlight Representational Similarity

To compare these neural response geometries to the structure in the intuitive similarity judgments, we calculated the Pearson’s correlation between the group-level intuitive similarity judgments and the neural RDM in each search sphere (**Figure 3b**). Correlations were considered significant for voxels that survived voxel-wise permutation tests (*p* < 0.01) and permutation-based cluster corrections (*q* < 0.05). We also repeated this process for each of the three model RDMs.

#### Three-Way Winner Map

To compare the whole-brain searchlight results across model RDMs, we calculated a 3-way winner map (**Figure 4**). In this map, we colored voxels according to the model RDM with the highest positive correlation with the neural RDM centered on that voxel. When there was a tie for the highest correlation, voxels were colored grey to indicate a lack of preference. In addition, the voxels’ saturation reflects the difference between the highest and next-highest correlations: voxels where one model RDM clearly dominated the others are colored more deeply. This analysis reflects an exploratory visualization using group-level data. As such, no statistics were done over this map.

## ACKNOWLEDGEMENTS

Funding for this project was provided by NIH grant S10OD020039 to Harvard University Center for Brain Science, NSF grant DGE1144152 to L.T., and the Star Family Challenge Grant to T.K. Thank you also to the Cambridge Writing Group and the Vision Lab Writing Club for their support during the writing process.

## Supplementary Information

### 1 Extended Methods

#### 1.1 Sociality and Interaction Envelope

In previous work Tarhan and Konkle (2020b), we found evidence that the visual cortex’s responses to action videos are organized by two features: *sociality* (whether an action is directed at a person) and the size of the *interaction envelope* (the spatial extent of an action’s effect on the world). The fMRI experiment in that work used the same set of action videos as in the current study. Because these features predicted action processing well in a large swathe of the brain, we wondered whether they captured information that informs the downstream processing supporting intuitive action understanding. To investigate this, we asked how well sociality and interaction envelope size could predict the intuitive similarity judgments.

To calculate the sociality and interaction envelope features for each video, we returned to the analysis that revealed this pattern in our previous work. In that work, a clustering analysis revealed 5 networks of brain regions. We interpreted the tuning of each network by examining the networks’ tuning to the different body parts involved in the action videos and the actions’ targets (what they were directed at, such as an object or a person). One network was tuned to actions directed at people, such as talking (sociality; **Supplemental Figure 2a**), while the remaining four were tuned to different interaction envelope sizes (**Supplemental Figure 2b**). For example, one network was tuned to small interaction envelopes, as in actions that involve fine, object-directed hand movements like knitting. At the other extreme, another network was tuned to larger interaction envelopes, as in actions that involved large movements of the whole body, directed at distant locations like a soccer penalty shot.

For each action video, we had measured which body parts were engaged by the action and what the action was directed at, using human ratings (see Tarhan and Konkle, 2020b for details). And for each network, we had calculated a tuning profile to these body parts and targets. So, to measure where each video fell along these five sociality and interaction envelope dimensions, we multiplied these body part and target ratings by each network’s tuning profile. This produced an estimate of how well each video aligned with each network’s preferred tuning. For example, a video of two people shaking hands was rated as being directed at other people and involved only the hands and arms. This video would then have a high value on the “sociality” and “small interaction envelope” dimensions, but lower values for the “medium interaction envelope,” “medium-large interaction envelope,” and “large interaction envelope” dimensions.

Finally, we assessed how well these features predict the intuitive similarity judgments in both experiments. In all three experiments, the sociality-interaction envelope features predicted the intuitive similarity judgments moderately well (**Supplemental Figure 2c**; mean cross-validated τ-a = 0.21, sd = 0.16 (Experiment 1); 0.26, s.d. = 0.16 (Experiment 2). However, these features performed worse than the three model RDMs based on the actors’ goals, movements, and the videos’ visual appearance. This suggests that, while sociality and interaction envelope predict action responses well in the visual cortex, they do not add much when predicting intuitive judgments.

## Supplemental Figures

**Supplemental Figure 1:**
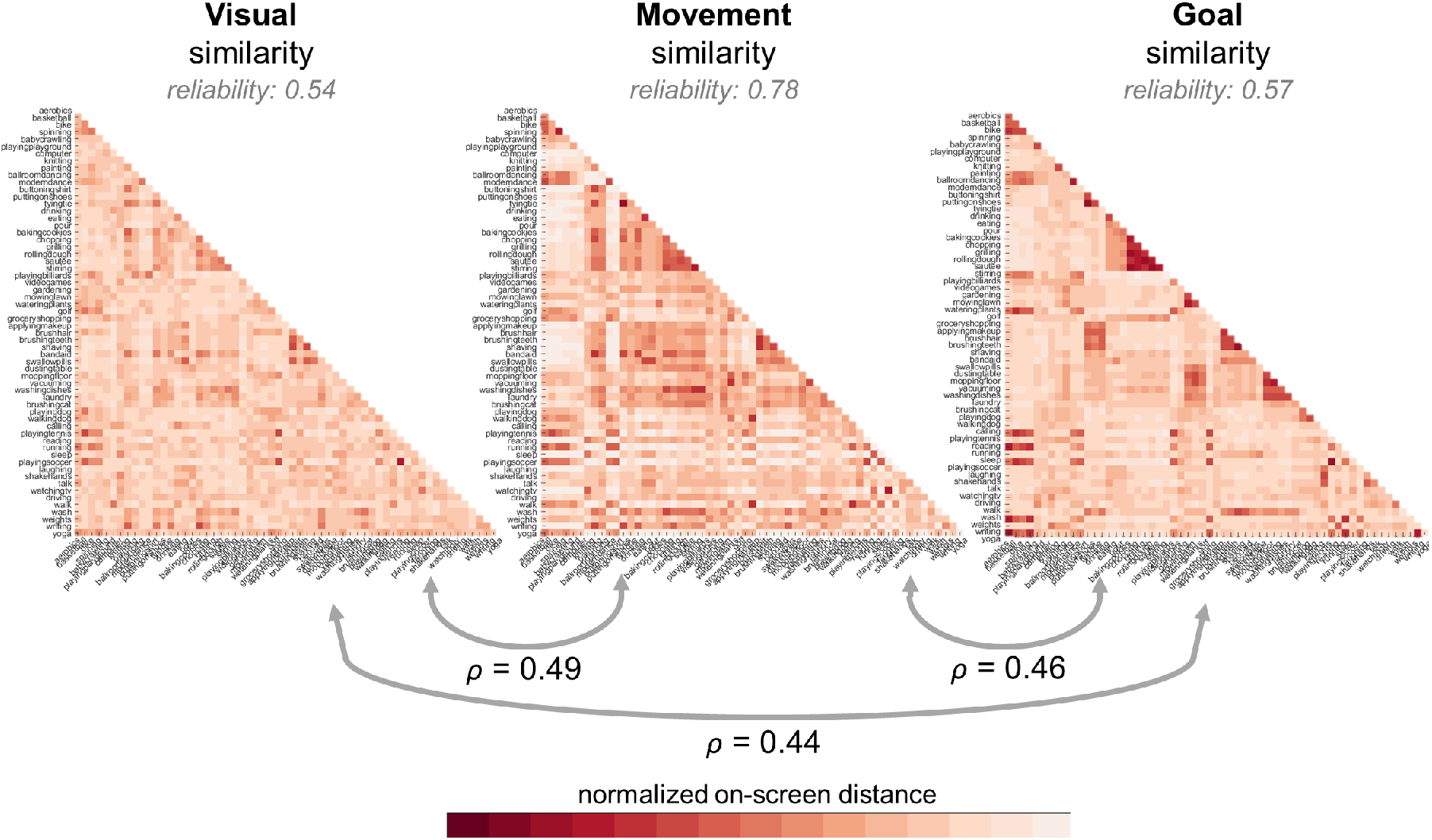
Comparing Model RDMs. Group-level Representational Dissimilarity Matrices (RDMs) are shown for the three types of action similarity judgments. Split-half reliabilities (after Spearman-Brown prophecy corrections) are listed for each type of judgment. Numbers beneath the grey arrows indicate the Spearmans’s correlations between these model RDMs.

**Supplemental Figure 2:**
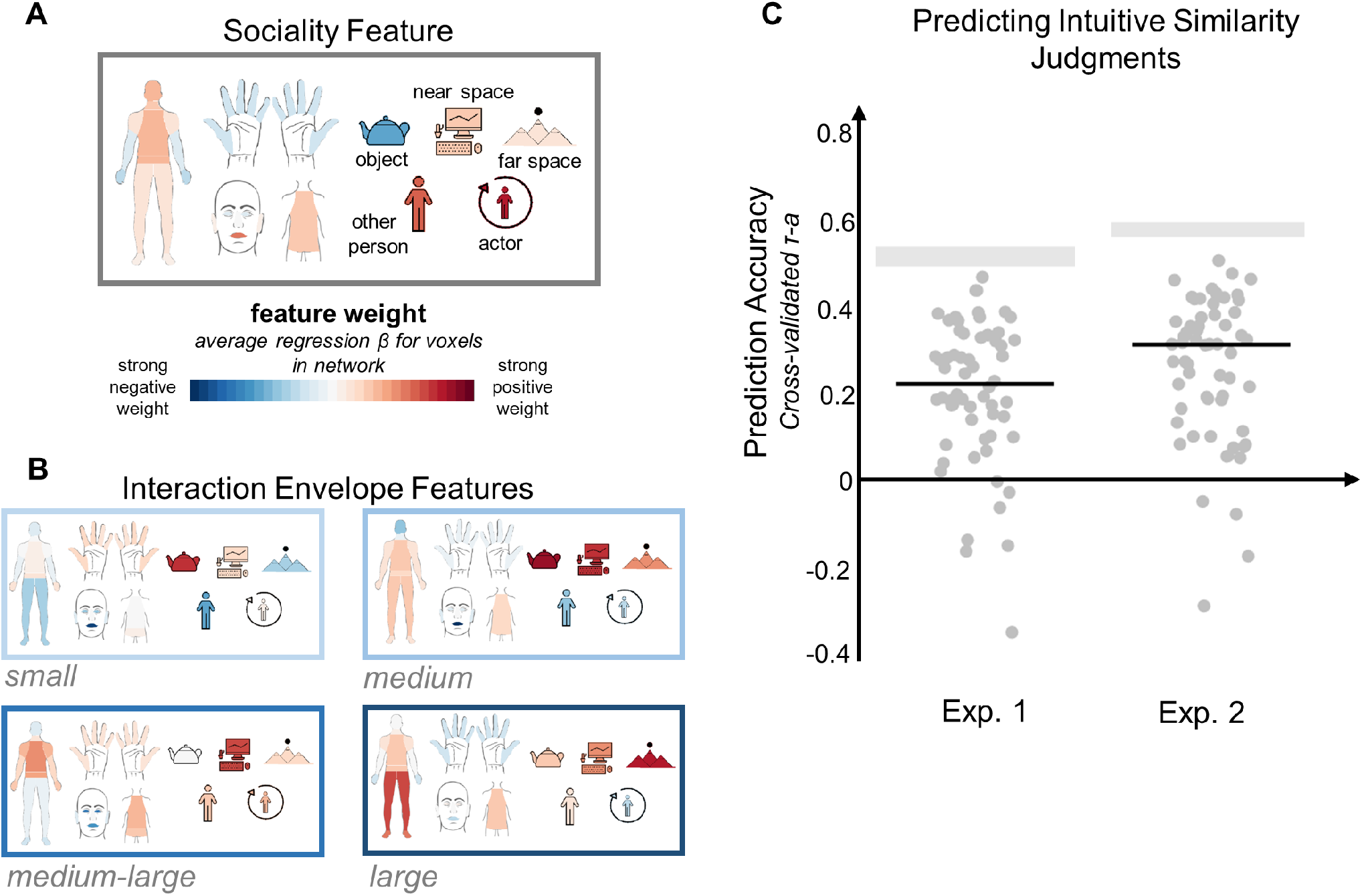
Sociality and Interaction Envelope. In prior work (Tarhan and Konkle, 2020b), we found that sociality and interaction envelope size predict action responses throughout the visual cortex. To assess whether these properties also influence downstream intuitive action processing, we asked how well similarity in actions’ sociality and the size of their interaction envelopes could predict intuitive similarity judgments. (A) Illustration of the sociality feature dimension, showing how much voxels in a right-lateralized network of brain regions were tuned to the body parts and targets involved in an action (Tarhan and Konkle, 2020b). Each body part and target is colored according to the strength of the network’s tuning – for example, this network was strongly tuned to actions directed at the actor or another person, but was not strongly tuned to actions directed at far space. (B) Illustrations of the four interaction envelope feature dimensions, which range from small envelopes around fine movements directed at objects (e.g., knitting) to large envelopes around coarser movements directed at distant locations (e.g., a soccer penalty shot). (C) We used these five dimensions – sociality and four sizes of interaction envelope – to predict intuitive similarity judgments about the actions. Prediction performance is plotted for the intuitive judgments measured in each experiment. Grey bars indicate the noise ceiling for each experiment. Horizontal black lines indicate the median prediction performance for each experiment, and grey dots plot performance on each iteration of the leave-1-condition-out procedure.

